# Multimodal Dynamics of Mental Fatigue and Their Selective Modulation by Acute Exercise: Effects on Memory and Creativity

**DOI:** 10.64898/2026.02.25.708006

**Authors:** Juliette Gélébart, Guillaume Digonet, Thomas Jacquet, Célia Ruffino, Ursula Debarnot

## Abstract

Mental fatigue (MF) arises from sustained cognitive load and produces a multisystem signature spanning subjective experience, task performance, cortical oscillations, and oculomotor dynamics. It may alter higher-order cognitive functions essential to everyday life, underscoring the need for preventive strategies. Although moderate aerobic exercise (EXO) facilitates recovery from MF, its influence on the onset and expression of MF when performed beforehand remains unexplored. This study provided a multimodal characterization of MF, assessed its impact on associative memory and divergent creativity, and examined whether prior EXO modulated these outcomes. Twenty-nine participants completed either 15 min of EXO or rest before a 35-min MF-inducing Time Load Dual Back task. Subjective fatigue and effort, performance, EEG activity, and eye-blink rate were continuously recorded; associative memory and divergent creativity were assessed pre-intervention and post-MF. Both groups showed progressive increases in MF and effort from 7 min onward, stable performance, and a rise in parieto-central alpha power at 18 min. The EXO group exhibited higher frontal-medial theta power and stable blink rates, whereas blink rate in REST increased at 21 min. EXO did not prevent subjective MF nor influence behavioral stability but modulated neurophysiological markers potentially related to compensatory control and dopaminergic regulation. Associative memory remained preserved in both groups, whereas creative flexibility increased in REST but not EXO, suggesting MF-related disinhibition in the former and preserved inhibitory control in the latter. These findings refine temporal and multimodal profile of MF and highlight the need to optimize exercise parameters and task demands to enhance preventive efficacy and guide interventions.

## 1. Introduction

Mental fatigue (MF), a psychobiological state induced by prolonged and/or intense cognitive activity, is characterized by subjective feelings of exhaustion, reduced motivation, impaired performance, and changes in brain activity (Tran et al., 2020). When sustained, it detrimentally affects productivity, decision-making, and overall cognitive efficiency (Pessiglione et al., 2025). Despite its increasing prevalence in daily life, the mechanisms underlying MF, as well as effective strategies to preserve mental performance and delay fatigue onset, remain insufficiently characterized.

Experimentally, MF can be induced through cognitively demanding tasks requiring sustained attention, vigilance, and inhibitory control such as the Time Load Dual Back (TLDB) or Stroop tasks (Goodman et al., 2025). Its detection can be captured across multiple complementary dimensions (Schampheleer et al., 2025). Subjectively, increased perceived exhaustion and effort induced by MF is typically assessed using Visual Analogue Scale (VAS ; Schampheleer et al., 2025). Behaviorally, it is reflected by slower reaction times, reduced accuracy, and higher error rates indicating a progressive decline in cognitive efficiency across time on task (Borragán et al., 2016; Kunasegaran et al., 2023). At the neurophysiological level, electroencephalography (EEG) studies show consistent temporal dynamics in brain oscillations during MF-inducing tasks (Tran et al., 2020). Alpha power increases within the first 15 to 45 minutes of the task, particularly over parietal regions (Hamann & Carstengerdes, 2023; Trejo et al., 2015). In parallel, frontal-midline theta activity increases progressively across the MF-inducing task (Tran et al., 2020), reflecting sustained executive control and compensatory effort (Wascher et al., 2014), although the precise timing of its emergence varies across studies depending on task demands, individual differences, and measurement approaches (Hamann & Carstengerdes, 2023; Wascher et al., 2014). Additionally, oculomotor markers, such as eye-blink frequency, rise during a demanding cognitive tasks and correlate positively with dopaminergic activity (Bafna & Hansen, 2021; Eckstein et al., 2017). To date, few studies have simultaneously combined these subjective, behavioral, and neurophysiological markers within the same experimental paradigm, limiting a comprehensive understanding of when and how MF manifests across complementary dimensions.

Studies focusing on cognition have shown that MF alters attentional and executive control systems (Behrens et al., 2023), thereby impairing multiple higher-order cognitive functions critical for everyday life, such as associative memory and creativity. Associative memory is one of the most common forms of memory used in everyday life, particularly in social interactions and learning contexts (Hargis & Castel, 2017; Suzuki, 2008). It requires the binding of multiple elements supported by attentional (Miller & Unsworth, 2021) and executive processes (Liu & Chung, 2021) and can be assessed using a Verbal Paired Associates (VPA) task. Although attentional and executive mechanisms are known to deteriorate under MF, the direct impact of experimentally induced MF on associative memory has not yet been tested. Creativity may also be influenced by MF, with potentially opposing effects depending on the type of creative process involved. Convergent thinking, which depends on executive controlled processing, typically declines under fatigue, whereas divergent thinking may paradoxically benefit from reduced inhibitory control, facilitating broader associative activation (Cassotti et al., 2016; Radel et al., 2015). This dissociation makes divergent creativity particularly valuable for characterizing the full spectrum of MF’s cognitive aftereffects. Thus, examining both associative memory and creativity within the same paradigm provides a more integrative understanding of the multifaceted cognitive consequences of MF.

Physical exercise has recently gained attention as a promising approach to mitigate the adverse effects of MF. Acute moderate aerobic exercise (EXO) performed *after* demanding cognitive tasks have been shown to reduce subjective fatigue and restore vigor (Jacquet, Poulin-Charronnat, Bard, Perra, et al., 2021; Oberste et al., 2021; Zhang et al., 2024). However, these studies have examined EXO exclusively as a recovery tool, leaving open the question of whether it could act as a preventive intervention when performed *before* exposure to a MF task. Moreover, most investigations have relied solely on self-reported fatigue, without jointly assessing behavioral, neural, and physiological markers (Díaz-García et al., 2021), limiting mechanistic insight. Nonetheless, there is strong theoretical support for the idea that performing exercise prior to a demanding cognitive task could delay or attenuate the onset of MF. Acute moderate exercise (EXO) enhances arousal and executive efficiency (Dupuy et al., 2024), increases prefrontal activation and dopaminergic activity (Basso & Suzuki, 2017; Byun et al., 2014), and counteract the alpha/theta synchronization and vigilance decline characteristic of MF (Tran et al., 2020). Beyond these immediate neural benefits, EXO also boosts brain-derived neurotrophic factor and modulates hippocampal–prefrontal functioning, which may help preserve memory and creative performance under cognitive strain (Endo et al., 2013; Knaepen et al., 2010). Taken together, these findings point to EXO as a potential proactive modulator of MF, capable of influencing its neurophysiological onset and subsequent cognitive effects.

Therefore, the present study aimed to characterize *when* and *how* MF manifests using a multimodal approach combining subjective, behavioral, electrophysiological and oculomotor markers, and to examine its aftereffects on higher-order cognition. A second objective was to determine whether performing an EXO before a fatiguing task could attenuate or delay the emergence of these MF markers. To this end, we quantified subjective fatigue and effort, task performance, EEG alpha and theta dynamics, and eye-blink rate during a 35-min Time Load Dual Back (TLDB) task, following either EXO or a REST intervention. We also assessed the aftereffects of MF on associative memory and divergent creativity. We expected MF to elicit progressive increases in subjective ratings, a gradual impairment in behavioral performance, a rise in alpha and theta power with time-on-task, and an elevation in blink rate, alongside impaired associative memory but enhanced divergent thinking. Regarding the effect of EXO, we hypothesized that, compared to REST, performing exercise prior to the TLDB task would yield lower subjective ratings, more stable behavioral performance, delayed increases in alpha and theta activity, and stable blink rates throughout the task, and would mitigate MF-related alterations in associative memory and creativity.

## 2. Materials and Methods

### 2.1. Participants

The sample size was determined using G*Power software (version 3.1.9.7) based on an a priori power analysis. For comparing two independent groups across multiple time points, an effect size f of 0.25 was used, representing a medium effect (Cohen, 1988). This effect size is consistent with previous studies examining MF interventions using comparable subjective and behavioral outcomes (Zhang et al., 2024). Statistical power and alpha level were set at 0.80 and 0.05, respectively. This yielded an estimated sample size of 24 participants (12 per group). Thus, thirty French-speaking participants with normal or corrected-to-normal vision and no history of neurological disease were included. Exclusion criteria included: (i) sedentary lifestyle or high-level athletic status, as determined by scores on the Global Physical Activity Questionnaire (GPAQ; scores outside the 400–4000 range; Bull et al., 2020); (ii) pathological boredom proneness, assessed using the Short Boredom Proneness Scale (Struk et al., 2017); and (iii) use of medication that could affect alertness, attention, or memory. Participants were instructed to sleep at least 7 hours the night before the experiment, refrain from alcohol, and vigorous physical activity the day priori, and refrain from caffeine and nicotine consumption before the experiment. Due to excessive movement artifacts that compromised EEG data quality, one participant was excluded from all analyses, resulting in a final sample of twenty-nine participants (mean age ± SD: 25.6 ± 5.0 years, 15 women). The present research was conducted in accordance with ethical guidelines and complies with the Declaration of Helsinki and all participants provided written informed consent prior to participation.

### 2.2. General procedure

The experiment was conducted in the morning, from 8.30 am to 11.00 am. Upon arrival, participants completed the Saint Mary’s Hospital questionnaire to assess their prior sleep quality. Then, participants provided baseline self-reports measures assessing their MF, mental effort (Visual Analogue Scale, VAS), alertness (Stanford Sleepiness Scale; Hoddes et al., 1972) and mood (Profile of Mood States ; Cayrou et al., 2003). After these assessments, participants performed the Verbal Paired Associates (VPA) task and the Alternative Uses Task (AUT; Guilford 1967), constituting the PRE-INTERVENTION session to assess respectively associative memory and divergent creativity performances. Participants were randomly allocated to one of two experimental groups (INTERVENTION phase). The exercise group (EXO), composed of 15 participants (24.6 ± 3.8 years; 7 women) engaged in 15 minutes of moderate-intensity cycling ergometer, and the rest-control group (REST), composed of 14 participants (26.8 ± 6.1 years, 10 women), remained seated while listening to a 15-minute neutral podcast. Following the INTERVENTION, participants’ EEG activity (32 electrodes) was recorded during a 5-minute resting-state period (RS-pre), during which they were asked to fixate on a black screen, eyes open, relax, and refrain from engaging in any specific thoughts. Then, participants performed the TLDB task for 35 minutes. During this task, EEG, oculomotor activity, and task performance were recorded, and subjective level of MF and mental effort were assessed every two breaks (i.e., every 7 minutes) using a VAS. Immediately following the TLDB task, a second 5-minute resting-state EEG was recorded (RS-post) under the same conditions as the RS-pre. Participants then reassessed their subjective MF, mental effort, alertness and mood. Finally, they performed again the VPA task and the AUT, during the POST-INTERVENTION session.

### 2.3. Physical exercise (EXO) and rest-control (REST) interventions

The EXO intervention consisted of 15 minutes of moderate-intensity cycling, during which participants maintained their heart rate between 50% and 60% of their maximum heart rate, monitored in real time using a Polar H10 chest strap (Polar Electro Oy, Kempele, Finland). Maximal heart rate was estimated using the validated age-based equation (Tanaka et al., 2001): HRmax = 208 – 0.7 × age.

In the REST group, participants remained seated and listened to a 15-minute neutral podcast on sneaker manufacturing. To minimize distractions, all participants faced a white wall throughout their respective conditions. At the end of the intervention, all participants rated their physical fatigue using the French translation of the Rating-of-Fatigue scale (ROF; 0 = “Not fatigued at all” to 10 = “Total fatigue and exhaustion – nothing left”; Brownstein et al., 2021). Participants in the EXO group reported an average fatigue score of 3.3 ± 1.5, while those in the REST group reported a score of 1.2 ± 0.4.

### 2.4. The Time Load Dual Back Task

The TLDB is a dual-task paradigm combining a classic 1-back working memory task with a concurrent interference task (Borragán et al., 2016). A continuous stream of letters and numbers was presented at a rate of one stimulus per second. For number appeared (1, 2, 3, 4, 6, 7, 8, or 9), participants pressed the “P” key if it the number was even and the “Z” key if it was odd. For letters (A, C, T, L, N, E, U, or P), participants pressed the spacebar if the letter matched the one immediately preceding it. Stimuli were displayed Arial, size 60, at center of a 16-inch screen (60 Hz refresh rate), and the task was implemented in Visual Studio Code, which also recorded accuracy and reaction times. The task consists of 10 blocks of 90 stimuli, each, separated by 30-second breaks, for a total duration of 34 minutes and 30 seconds. Performance was analyzed in ten consecutive 3-minute time bins (T1 to T10), capturing the evolution of accuracy and reaction time over the course of the task (e.g., T1: 0-3 min; T2: 3.5-6.5 min, […], T10: 31.5-34.5 min).

### 2.5. Subjective assessments of alertness, mood, mental fatigue and effort

At PRE- and POST-INTERVENTION participants rated their level of alertness using the Stanford Sleepiness Scale (SSS; Hoddes et al., 1972), a single-item scale consisting of seven descriptive statements ranging from 1 (‘Feeling active and vital; alert, wide awake’) to 7 (‘No longer fighting sleep, sleep onset soon; having dream-like thoughts’). Mood was assessed using the 37-item POMS-SF (Cayrou et al., 2003), rated on a 5-point scale and comprising six subscales (Depression, Anxiety, Anger, Fatigue, Vigor, Confusion), with high internal consistency (α = 0.80–0.91). Subjective ratings of MF and perceived mental effort were collected at PRE-INTERVENTION, intermittently during the TLDB (t7, t14, t21, t28), and at POST-INTERVENTION using two separate 10-point visual analogue scales (VAS; 0 = “Not at all”, 10 = “Extremely”). Participants responded to the questions: *“How mentally fatigued do you feel right now?”* and *“How much mental effort did you exert during the task?”*

### 2.6. Neural markers of mental fatigue

#### 2.6.1. EEG recording and preprocessing

EEG was recorded using 32 electrodes positioned according to the international 0–20 system. Electrode impedance was kept below 10 kΩ. Eye blinks were monitored using an electrode placed below the left eye. Signals were sampled at 1000 Hz using BrainAmp amplifiers and recorded with BrainVision Recorder software (Brain Products®, Munich, Germany). Offline preprocessing was performed using the BrainVision® System (Brain Products®, Munich, Germany). Data were down-sampled to 256 Hz, re-referenced from the initial mastoid reference (TP9 and TP10) to the average reference, and filtered using a 1 Hz high-pass filter and a 50 Hz notch filter. Independent component analysis was applied to identify and remove components reflecting eye and muscle artifacts based on visual inspection. EEG data from the different conditions (RS-pre, time-on-task T1 to T10, and RS-post) were segmented into 5000 ms epochs for further analysis.

#### 2.6.2. EEG Spectral Analysis

Spectral analysis was conducted during the TLDB task and the two 5-minute resting state periods (RS-pre and RS-post). Fast Fourier Transforms (FFT) were used to compute the power of individual epochs in two frequency bands: theta (4-7 Hz) and alpha (8-12 Hz). To examine spatial changes in spectral power, nine regions of interest (ROIs) were determined by averaging signals across electrode clusters, as proposed by Arendsen et al. (2020): Anterior Left (Ant_L: FP1, AF3, F3, F7, FC5), Anterior Median (Ant_M: Fz, FC1, FC2), Anterior Right (Ant_R: FP2, AF4, F4, F8, FC6), Central Left (Cen_L: C3, T7, CP5), Central Median (Cen_M: Cz, CP1, CP2), Central Right (Cen_R: C4, T8, CP6), Posterior Left (Post_L: P7, P3, PO3, O1), Posterior Median (Post_M: Pz, Oz), and Posterior Right (Post_R: P4, P8, PO4, O2).

#### 2.6.3. Blink rate analysis

Blink rate was estimated from the infraorbital electrode (below the left eye) and calculated as the average number of blinks per minute for each time-on-task interval (T1 to T10). To account for inter-individual variability and allow between-group comparisons, blink rate changes were expressed as a percentage relative to T1.

### 2.7. Verbal Paired Associates task

The VPA is a widely used instrument for testing associative memory and has been a continuous subtest within the Wechsler Memory Scale (WMS-IV; Wechsler, 2009). For this task, participants were instructed to memorize a list of 24 cue–response word pairs, each displayed one by one on the screen for 4 seconds. They were then asked to recall and type the response word associated with each cue. The task included three recall trials, during which the same 24 pairs were presented in a randomized order. The task was administrated at PRE- and POST-INTERVENTION (i.e., after the TLDB task), using different sets of 24 word pairs each time. All the words were French concrete noun objects of two or three syllables. Half of the pairs were semantically related (e.g., pencil-eraser) and half were unrelated (e.g., window-coffee). Words were matched for familiarity and imageability using French psycholinguistic norms from (Bonin et al., 2003). For each participant, the proportion of correctly recalled words was computed for each trial and session (PRE-INTERVENTION, POST-INTERVENTION), then averaged by group (EXO, REST).

### 2.8. Alternative Uses Task (AUT)

The AUT was used to assess divergent thinking. Participants were instructed to generate as many uses as alternative uses as possible for three common household items, excluding their typical function. They had two minutes per object to type their responses on the computer. Two lists of items were used (list 1: bucket, shoe, newspaper; list 2: umbrella, can, paperclip), with the order counterbalanced across participants and across PRE- and POST-INTERVENTION. During a brief familiarization phase, participants practiced with “pen” and “padlock”. Creativity was assessed using four standard components derived from a consensual assessment approach (Silvia et al., 2009). *Fluency* referred to the number of uses generated per item. *Flexibility was calculated* as the number of distinct usage categories divided by the fluency score. *Elaboration* corresponded to the mean number of words per item, and *originality* measured the uniqueness of ideas to others. Scores obtained for each object were then averaged. Fluency and flexibility were scored by two raters to reduce subjectivity bias (Cronbach’s alpha was 1 for fluency, 0.812 for flexibility). Elaboration and originality were scored automatically using Large Language Models (LLMs) based on *supervised learning trained on previously scored responses* (Organisciak et al., 2023) with ratings on a 1-5 scale (1 = very unoriginal, 5 = very original). Scores were averaged across objects and compared between EXO and REST groups for each creativity component.

### 2.9. Statistical Analyses

To examine the impact of MF on alertness and mood, as well as the modulation induced by the intervention, repeated-measures ANOVAs (ANOVA_RM_) were conducted on the corresponding subjective scales. For the SSS, a two-way ANOVA_RM_ was performed with TIME (PRE-INTERVENTION, POST-INTERVENTION) as a within-subject factor and INTERVENTION (EXO, REST) as a between-subject factor. For the POMS, a three-way ANOVA_RM_ was conducted with DIMENSIONS (Anxiety, Depression, Anger, Vigor, Fatigue, Confusion) and TIME (PRE-INTERVENTION, POST-INTERVENTION) as within-subject factors, and INTERVENTION (EXO, REST) as a between-subject factor.

To assess the impact of MF on subjective and behavioral markers, as well as intervention-related modulation (EXO vs. REST), two-way ANOVA_RM_ were performed on MF and mental effort ratings (VAS), with PERIOD (PRE-INTERVENTION, t7, t14, t21, t28, POST-INTERVENTION) as a within-subject factor and INTERVENTION (EXO, REST) as a between-subject factor. Reaction times and accuracy during the TLDB were analyzed using ANOVA_RM_ with TIME-ON-TASK (T1 – T10) as a within-subject factor and INTERVENTION as a between-subject factor. To assess the neural markers of MF, theta and alpha band power were analyzed separately using ANOVA_RM_. For EEG recordings during the TLDB task, ANOVA_RM_ were performed on theta and alpha power, with TIME-ON-TASK (T1 to T10) and REGION (Ant_L, Ant_M, Ant_R, Cen_L, Cen_M, Cen_R, Post_L, Post_M, Post_R) as within-subject factors, and INTERVENTION as the between-subject factor. Blink rate was analyzed using a ANOVA_RM_ with TIME-ON-TASK (T1 to T10) as the within-subject factor and INTERVENTION as the between-subject factor.

To examine the impact of MF and whether exercise modulated post-fatigue cognitive functioning, ANOVA_RM_ were conducted on the VPA task, namely, the number of correctly recalled words was analyzed using a three-way ANOVA_RM_ with TIME (PRE-INTERVENTION VS POST-INTERVENTION) and TRIAL (1, 2, 3) as within-subject factors, and INTERVENTION as the between-subject factor. For the AUT, two-way ANOVA_RM_ were conducted on creativity component (flexibility, fluency, elaboration and originality) with TIME as within-subject factors, and INTERVENTION as the between-subject factor.

To verify initial group equivalence, Mann-Whitney U tests were conducted to compare physical activity levels, sleep parameters (duration and quality), and vigilance levels (SSS) between the EXO and REST groups. For POMS scores, a one-way ANOVA was performed to compare mood states across the six dimensions at PRE-INTERVENTION.

All statistical analyses were performed using JASP software (version 0.16.1.0). Prior to conducting parametric tests, assumptions of normality and homogeneity of variance were assessed using Shapiro-Wilk tests and Levene’s tests, respectively. The significance threshold was set at *p* < .05 (two-tailed). When the assumption of sphericity was violated, Greenhouse–Geisser corrections were applied. To control for multiple comparisons, Bonferroni corrections were applied where appropriate including for post hoc tests following significant effects or interactions. Partial eta-squared (η^2^p) values were reported as measures of effect size.

## 3. Results

No significant baseline differences were observed between groups in physical activity (U = 73.50, *p* = 0.666), sleep parameters (duration: U = 73.50, *p* = 0.267, quality: U = 125, p=0.195), vigilance levels (SSS; U = 93, *p* = 0.817), and POMS scores (F_(1,28)_ = 0.469, *p* = 0.500, η_p_^2^ = 0.018) at PRE-INTERVENTION indicating initial homogeneity between the two experimental groups (see Supplementary Table S1).

### 3.1. Subjective fatigue increased while behavioral performances remained stable, with no effect of EXO

Vigilance (SSS) decreased from the PRE- to POST-INTERVENTION (main effect of TIME; F_(1,28)_ = 18.331, *p* < 0.001, η _p;_^2^ = 0.414), with no group differences or time × intervention interaction (all *ps* > 0.23, and all η _p_^2^ < 0.055), indicating that EXO did not modulate changes in sleepiness (Fig. 2A). Mood states (POMS) similarly changed over time (dimension × time interaction; F^(2377, 59.424)^ = 16.288, *p* < 0.001, η _p_^2^ = 0.394) driven by decreased VIG (t(28) = 5.949, *p* < 0.001) and increased FAT (t(28) = -6.596, *p* < 0.001). Again, no main effect of intervention was found (F_(1,28)_ = 1.030, *p* = 0.320, η _p_^2^ = 0.040), suggesting that these mood changes occurred independently of EXO (Fig. 2B).

**Figure 1.**
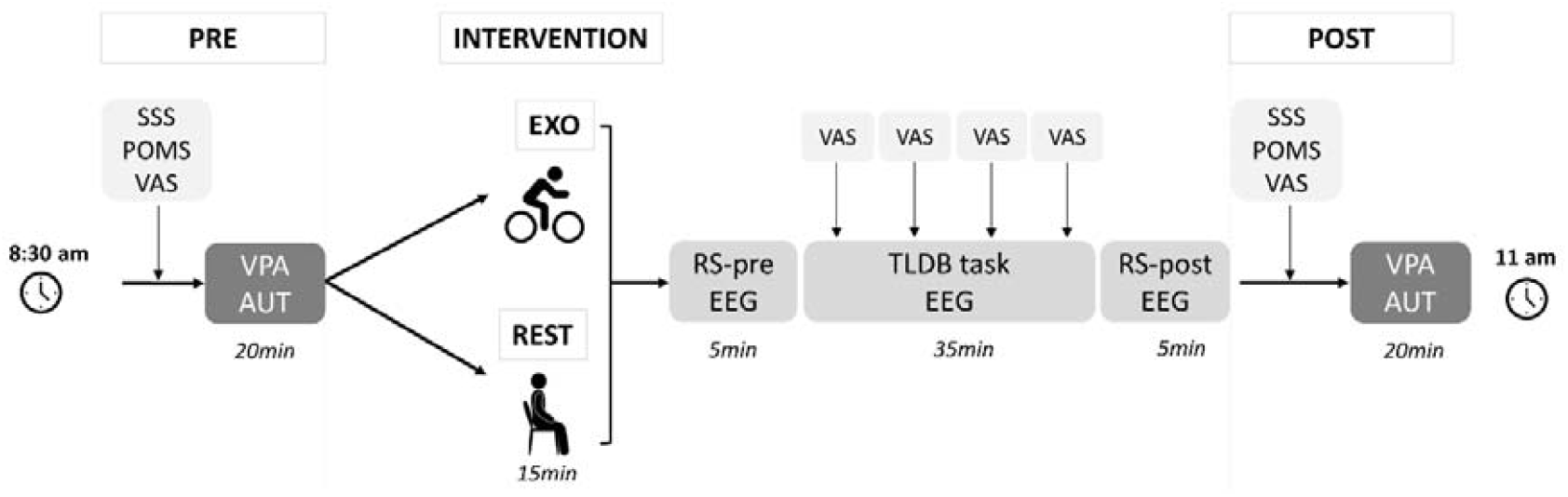
Overview of the experimental protocol. Participants completed pre- and post-intervention questionnaires, (SSS: Stanford Sleepiness Scale; POMS: Profile of Mood States scale; VAS: mental fatigue and effort visual analogue scale), followed by Verbal Paired Associates task (VPA) and Alternative Uses Task (AUT). Between assessments, they were assigned to moderate exercise (EXO) or rest (REST), and then performed the Time Load Dual Back task (TLDB) under EEG with repeated fatigue/effort ratings.

**Figure 2:**
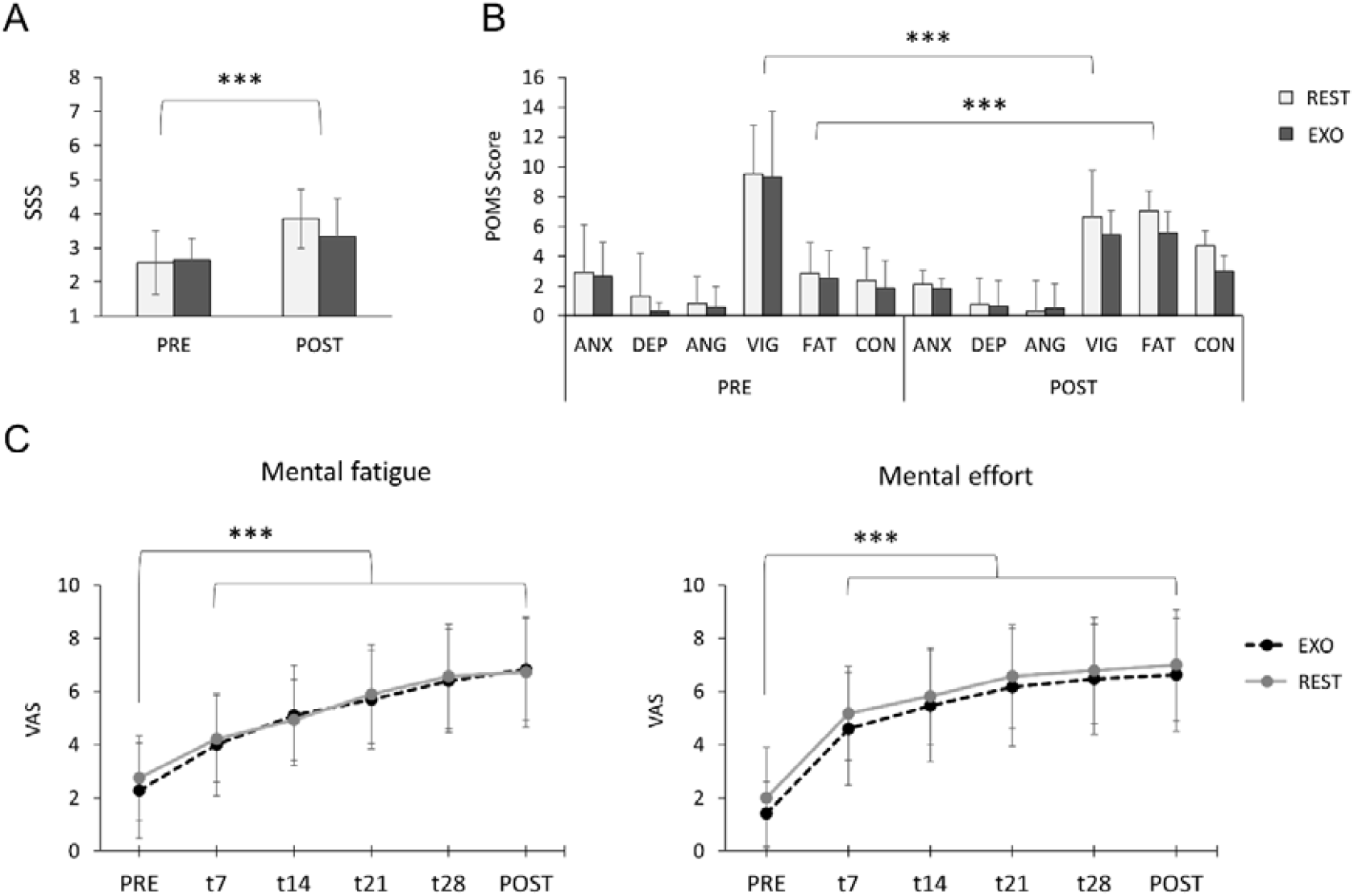
**A**. vigilance scores on the Stanford Sleepiness Scale (SSS) according to time and intervention. **B**. Profile of Mood State (POMS) scores according to the six dimensions, time and intervention. **C**. Subjective ratings of mental fatigue (left) and mental effort (right) according to the period and intervention. ANX, anxiety-tension; DEP, depression; ANG, anger-hostility; VIG, vigor-activity; FAT, fatigue-inertia; CON, confusion-bewilderment. *** p < 0.001.

During the TLDB, subjective MF and perceived effort increased progressively from the first rating (7 min), as shown by a main effect of the period both for MF (F_(2.379,61.856)_ = 63.262, *p* < 0.001, η _p_^2^ = 0.709) and effort (F_(2.452,61.296)_ = 55.668, *p* < 0.001, η_p_^2^ = 0.690). However, EXO did not modify the evolution of these measures: neither a main effect of intervention nor a period × intervention interaction emerged (all *ps* > 0.60 and η_p_^2^ < 0.016; Fig. 2C).

The mean reaction time did not change with time-on-task (F_(3.481, 80.061)_ = 2.398, *p* = 0.065, η_p_^2^ = 0.052), and no main effect of intervention (F_(1,28)_ = 0.011, *p* = 0.918, η_p_^2^ = 0.002), or time-on-task x intervention interaction (F_(3.481, 80.061)_ = 2.525, *p* = 0.055, η_p_^2^ = 0.049) emerged. Accuracy showed the same pattern, with no effect of time-on-task (F_(9,207)_ = 0.755, *p* = 0.658, η_p_^2^ = 0.032), intervention (F_(1,28)_ = 0.092, *p* = 0.765, η_p_^2^ = 0.004), nor their interaction (F_(9,207)_ = 0.319, *p* = 0.968, η_p_^2^ = 0.014). Together these results indicate that both reaction time and accuracy remained stable throughout the TLDB task regardless of the intervention.

### 3.2. Neural markers of MF: alpha increased with time-on-task, while EXO enhanced frontal theta

Alpha showed a clear temporal signature of MF with a main effect of TIME-ON-TASK emerged across several regions, increasing from T6 onward (18 minutes) compared to T1 in Central (Cen_L, Cen_R), Posterior (Post_L, Post_R) regions (all *ps* > 0.05; Fig. 3A). Importantly, EXO did not modify alpha dynamics, as neither a main effect of intervention nor an interaction with time-on-task was observed (all *ps* > 0.31). Theta activity, in contrast, did not display a time-dependent MF signature, with no main effect of TIME-ON-TASK (F_(4.004,88.090)_ = 1.597, *p* = 0.182, η_p_^2^ = 0.068). However, theta was influenced by EXO: a main effect of intervention (F_(1,28)_ = 4.480, *p* = 0.046, η_p_^2^ = 0.169), indicated consistently higher frontal-medial theta power (Ant_M) in the EXO group (all *ps* < 0.001 ; Fig. 3B). No significant effect of, or any significant interactions were observed (all *ps* > 0.05).

**Figure 3.**
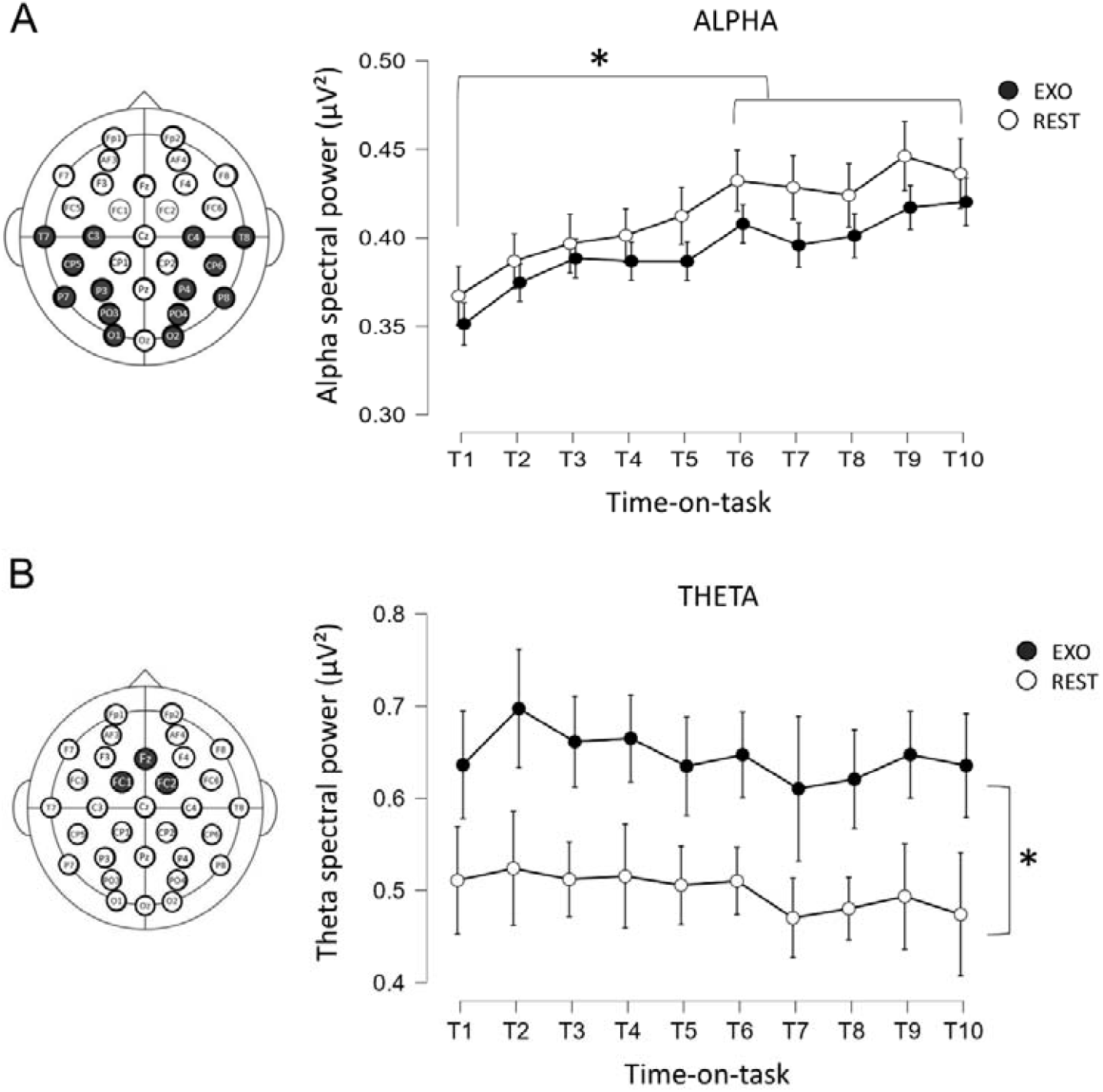
Evolution of alpha and theta power during the TLDB task, according to the intervention. **A.** Alpha power averaged across central and posterior electrodes for the EXO and REST groups as a function of time-on-task (T1-T10). **B**. Theta power averaged across anterior-medial electrodes for the EXO and REST groups as a function of time-on-task. The topographic maps on the left highlight the scalp regions where significant power changes were observed. The error bars represent the standard error of the mean. *p < 0.05.

Analysis of the percentage change relative to T1 revealed a significant time-on-task × intervention interaction (F_(2.515, 63.341)_ = 2.887, *p* = 0.045, η_p_^2^ = 0.116; see Fig. 4), indicating differential evolution across groups. In the REST group, blink rate increased from T7 onward (all *ps* < 0.003), reaching a +74 % ± 23 rise at T10. In contrast, the EXO group maintained a stable blink rate across the entire task, with only a minimal change at T10 (+ 9 % ± 11%). The mean blink rate across the full task did not differ between groups (EXO: 19.7 ± 8.9; REST: 19.6 ± 9.3 blinks/min).

**Figure 4.**
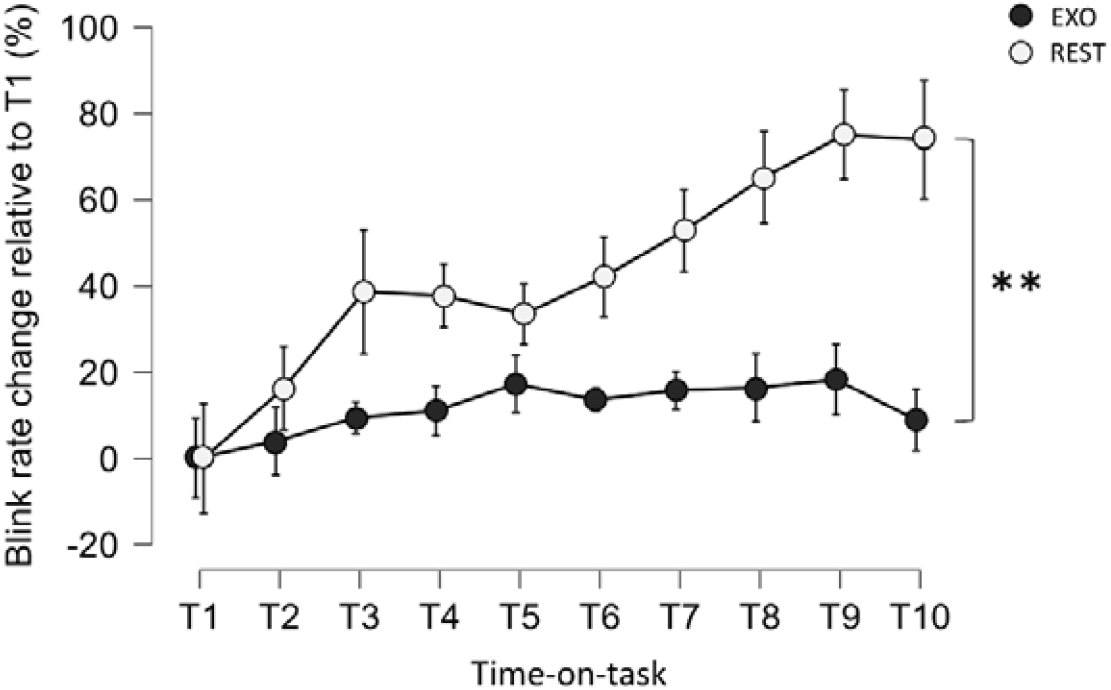
Evolution of blink rate as a function of time-on-task, expressed as a percentage relative to the blink rate at T1. The error bars represent the standard error of the mean. **p < 0.01.

**Figure 5.**
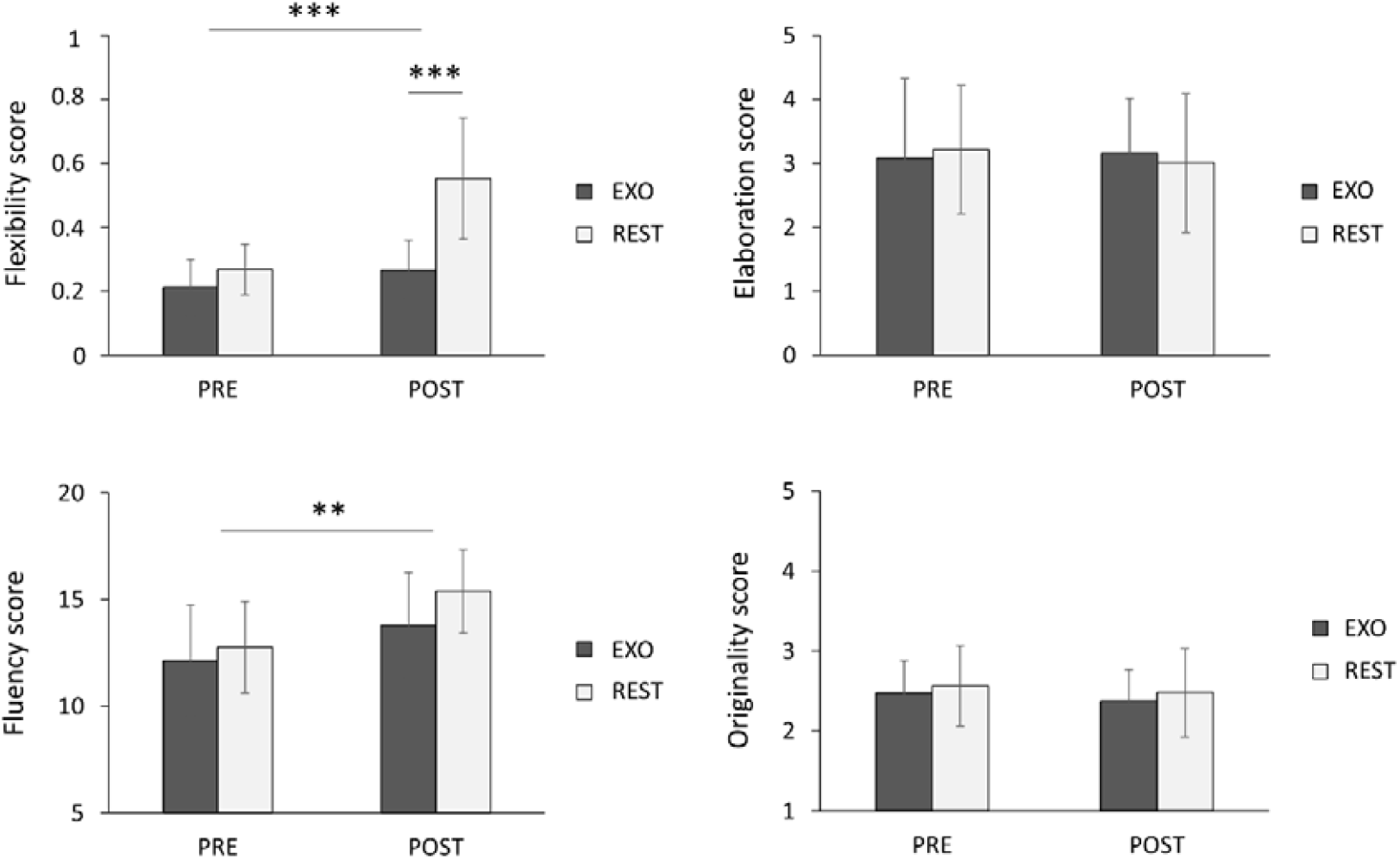
Means of AUT scores assessing divergent creativity obtained before and after the mental fatigue induction task in both groups. **p < 0.01, ***p < 0.001.

### 3.3. MF spares memory while selectively enhancing divergent creativity, an effect modulated by exercise

Associative memory performance (VPA) remained high in both groups, with near-ceiling accuracy by the third recall trial at both PRE- and POST-INTERVENTION (≥ 94%). A main TRIAL effect confirmed improved recall across repetitions (F_(1.222, 27.209)_ = 123.582, *p* < 0.001, η_p_^2^ = 0.867), but no effects of time, intervention or their interactions were observed (all *ps* > 0.35, η_p_^2^ < 0.075). Thus, MF did not degrade associative memory performance, and EXO did not modulate memory outcomes.

For the AUT, flexibility scores, reflecting the ability to shift between conceptual categories, revealed a main effect of intervention (F_(1,28)_ = 27.880, *p* < 0.001, ηp^2^ = 0.517), time (F_(1,28)_ = 27.827, *p* < 0.001, η_p_^2^ = 0.517), and a intervention x time interaction (F_(1,28)_ = 12.803, *p* = 0.001, η_p_^2^ = 0.330). REST participants displayed a marked increase in flexibility from PRE to POST-INTERVENTION (t(28) = -6.260, *p* < 0.001), but no improvement emerged in the EXO group (t(28) = -1.200, *p* > 0.05 ; Fig. 4). Fluency, indexing the quantity of generated ideas, increased overall from PRE to POST (F_(1,28)_ = 11.756, *p* = 0.002, η_p_^2^ = 0.311), with no EXO effect. By contrast, elaboration, corresponding to the mean number of words per item, and originality, reflecting the uniqueness of ideas, remained unchanged, with no influence of time, intervention or their interaction (all *PS* > 0.32, η*p*^2^ < 0.035).

## 4. Discussion

This study characterized MF using multimodal subjective, behavioral, electrophysiological and oculomotor markers, examined its cognitive aftereffects, and tested whether EXO could mitigate or delay the onset of MF-related signatures. Across the TLDB task, subjective fatigue and perceived effort rose early (7 min), while behavioral performance remained stable. MF was marked by an increase in parieto-central alpha power around 18 minutes, whereas frontal-medial theta did not show a progressive time-on-task rise, although theta activity remained consistently higher in the EXO group than in REST. Oculomotor dynamics further differentiated the groups, with blink rate increasing in REST from 21 min onward while remaining stable after EXO. At the cognitive level, associative memory performance was preserved in both groups, whereas divergent thinking improved after the MF task, with a larger gain in flexibility in REST than in EXO.

### 4.1. Multimodal characterization of mental fatigue dynamics

As expected, subjective fatigue increased during the TLDB task in both groups. However, unlike the performance declines typically observed with an individually calibrated version of the task (Borragán et al., 2016, 2017), behavioral performance remained stable over time in our study. One explanation may lie in our fixed-difficulty TLDB, which, in contrast to adaptive paradigms, did not adjust task demands to individual capacity and may therefore have been less sensitive to subtle performance deterioration. Alternatively, stable accuracy despite rising subjective fatigue has been reported in prior MF studies (Jacquet, Poulin-Charronnat, Bard, & Lepers, 2021; Pageaux et al., 2015) and is generally interpreted as reflecting compensatory control mechanisms recruited to sustain performance under increasing cognitive load (Badarin et al., 2025; Hockey, 2011). Consistent with this interpretation, participants exhibited a progressive rise in perceived mental effort, suggesting that preserved performance likely reflects the mobilization of compensatory cognitive resources rather than the absence of fatigue.

At the neural level, alpha power increased around the 18^th^ minute, mainly over centro-posterior regions. This pattern replicates previous EEG findings showing MF-related alpha enhancement and aligns with the timing reported for fatigue-associated neural shifts (Boksem et al., 2005; Tran et al., 2020; Wascher et al., 2014). Such increases have been interpreted as markers of progressive attentional disengagement, with cortical inhibitory mechanisms suppressing irrelevant external input to preserve performance despite reduced vigilance (Bazanova & Vernon, 2014; Clayton et al., 2015). In contrast, fronto-medial theta power remained stable across the task. Given its established role as an index of sustained cognitive control (for a review, see Cavanagh & Frank, 2014), stable theta activity likely reflects the constant executive demands of the TLDB task (Xie et al., 2021). It is also plausible that the task duration was insufficient to reveal a gradual increase in theta power, as such changes are typically detected only in protocols extending beyond 60 to 90 minutes of continuous cognitive effort (Hamann & Carstengerdes, 2023; Wascher et al., 2014).

Oculomotor dynamics provided an additional marker of MF development. Blink rate increased progressively in the REST group from the 20^th^ minute onward, consistent with evidence that eye-blink frequency rises during sustained cognitive effort and has been associated with dopaminergic modulation (Maffei & Angrilli, 2018). Given that both hypo- and hyperdopaminergic states can impair cognitive control mechanisms, including working memory (Cools & D’Esposito, 2011), this increase may reflect fatigue-related dysregulation in dopaminergic function, potentially contributing to reduced efficiency of cognitive control processes.

### 4.2. Prior EXO influences neural, not experiential dimensions of MF

EXO did not mitigate subjective MF, nor did it change behavioral performance relative to REST, or delay the increase in alpha power. However, clear EXO-related differences emerged in theta activity, with the EXO group exhibiting consistently higher frontal-medial theta power than the REST group throughout the task. This pattern is consistent with recent evidence showing that acute moderate exercise potentiates frontal theta during subsequent executive tasks (Griggs et al., 2023) and may indicate enhanced responsiveness of cognitive control systems to the sustained demands imposed by the TLDB task (Cavanagh & Frank, 2014). However, this heightened neural engagement did not translate into observable behavioral improvements during the task, consistent with numerous studies showing that changes in neural activity may precede or occur independently of changes in behavioral performance (Neubauer & Fink, 2009).

Oculomotor activity also distinguished the two groups, with blink rate increased progressively in the REST condition, whereas it remained stable after EXO. This divergence suggests that MF may have induced a progressive dopaminergic depletion in the REST group (Bafna & Hansen, 2021; Eckstein et al., 2017). Given that acute moderate exercise increases dopamine release in the brain (Ando et al., 2024), EXO may have compensated for the MF-related dopaminergic depletion observed in the REST group, thereby maintaining dopaminergic tone and preserving the efficiency of cognitive control processes during prolonged effort (Cools & D’Esposito, 2011). Although this interpretation must be considered cautiously in the absence of direct neurochemical measurements in our study, the divergence in blink rate trajectories between groups provides preliminary evidence that EXO can mitigate at least some oculomotor manifestations of MF.

### 4.3. MF spares associative memory but promotes creative flexibility, an effect prevented by prior EXO

Associative memory was not affected by MF, as VPA performance remained stable from pre- to post-test. This absence of behavioral change is consistent with some studies showing that associative memory can remain intact under conditions of reduced attentional resources, particularly when encoding relies on pre-existing semantic relationships between items (Wong et al., 2021). In the present study, the use of semantically related word pairs may have favored the recruitment of more automatic encoding processes that are less dependent on the executive binding mechanisms vulnerable to MF (Miller & Unsworth, 2021; Tibon et al., 2014). The absence of exercise-related facilitation on associative memory in the EXO condition is consistent with evidence highlighting the critical role of timing in the acute exercise–memory relationship. Benefits of acute exercise are most reliably observed when encoding occurs within a narrow post-exercise window (Crawford & Loprinzi, 2019; Loprinzi et al., 2019). In the present study, designed to examine associative memory performance following a sustained MF-inducing task, encoding took place more than 35 minutes after EXO, placing the exercise bout in a temporal window that may have attenuated benefits on encoding.

In contrast, creative flexibility improved after the TLDB task in the REST group, whereas this facilitation was abolished for the EXO group. The REST pattern closely resembles findings by Radel et al. (2015) who showed that sustained engagement in demanding executive tasks reduces inhibitory control and temporarily promotes divergent thinking. Our data suggest a comparable mechanism whereby MF may have reduced inhibitory constraints, enabling broader associative activation and thereby enhancing creative flexibility. This behavioral pattern is further supported by the blink-rate profile of the REST group. Spontaneous blink rate is linked to dopaminergic functioning and follows an inverted-U relationship with cognitive flexibility (Chermahini and Hommel, 2010). The progressive blink-rate increase observed during the TLDB task is consistent with MF-related dopaminergic dysregulation (Maffei & Angrilli, 2018) and may have shifted participants toward a reduced inhibition conducive to divergent thinking (Radel et al., 2015). By contrast, given that acute exercise can enhance inhibitory control (Levin et al., 2021) and modulate dopaminergic neurotransmission (Heijnen et al., 2016; Marques et al., 2021), it is plausible that EXO helped preserve inhibitory processes and prevented the transient disinhibition induced by MF alone. However, these interpretations remain preliminary and should be tested directly in future studies designed to manipulate inhibitory demands and dopaminergic functioning explicitly.

## 5. Conclusion

Our study provides converging multimodal evidence that MF manifests rapidly across subjective, electrophysiological, and oculomotor dimensions, even when behavioral performance remains preserved, reflecting compensatory control, attentional disengagement, and likely dopaminergic dysregulation. Acute moderate exercise performed beforehand did not attenuate or delay these experiential signatures of MF but selectively modulated neurophysiological markers producing consistently higher frontal-medial theta power and stable blink rates throughout the fatiguing task. Regarding the aftereffects of MF on higher-order cognition, MF spared associative memory but transiently enhanced divergent creative flexibility suggesting that MF-induced reductions in inhibitory control may promote creative expansion while exercise helps maintain inhibitory regulation. Together, these findings dissociate experiential, behavioral, and neural expressions of MF and identify acute moderate exercise as a modulator of the state, but not the existence, of MF. They also highlight the need to clarify when and how exercise-induced neuromodulation can yield functional cognitive benefits in real-world contexts.

## CRediT authorship contribution statement

Juliette Gélébart: Writing — original draft, Visualization, Methodology, Investigation, Formal analysis, Data curation, Conceptualization. Guillaume Digonet: Writing — review & editing, Methodology, Software, Conceptualization. Thomas Jacquet: Writing — review & editing, Methodology, Conceptualization. Célia Ruffino: Writing — review & editing, Supervision, Methodology, Conceptualization. Ursula Debarnot: Writing — review & editing, Supervision, Funding acquisition, Conceptualization.

## Data availability

The raw EEG data are available from the corresponding author upon reasonable request.

